# Cocaine extended access gates compulsive-like seeking in rats

**DOI:** 10.1101/2025.02.07.637011

**Authors:** Mónica Tapia-Pacheco, Jeanne-Laure de Peretti, Lucie Vignal, Maya Williams, Christelle Baunez, Jean-Marc Goaillard, Mickael Degoulet

## Abstract

When does compulsive-like drug seeking emerge? Despite decades of research, and critical advances in our understanding of brain processes leading to addiction, this question remains widely unanswered. So far, behavioral models assessing compulsive-like cocaine seeking following an extended access to cocaine failed to capture the development of its compulsive seeking. In fact, compulsive-like animals immediately displayed pathological seeking when facing its negative consequences of drug seeking, usually materialized by an unescapable mild electric shock on the paws. Here, we designed a new task, ‘Punished Seeking during Extended access’ (PSE), by inserting punished seeking trials within the sessions of extended access to cocaine. We show that compulsive-like cocaine seeking progressively emerges after several PSE sessions, once cocaine intake has been escalated. We thus provide the addiction community a pertinent model to explore brain mechanisms sustaining the emergence of compulsive-like cocaine seeking.

## Main Text

The persistent inability to refrain harmful use of a substance is the most discriminative criterion for addiction diagnosis (*1*), leading to detrimental personal, social, and health damages for users. As in human, vulnerability to compulsively seek (*2–5*) or take drugs (*6–10*) is also observed in subsets of rodents. The expression of compulsive-like behaviors relies on corticostriatal circuits, notably implying dysregulations in prefrontal and orbitofrontal cortical areas that yield to an activity shift from the ventral to the dorsolateral striatum (*2, 11–13*), which likely affect goal-directed behaviors, decision-makings, and habit formations (*14, 15*).

This compulsive trait involves two distinct, but intermingled, behavioral actions: drug taking, which is mostly driven by the pharmacological effects of a substance, and drug seeking that involves complex foraging strategies to obtain the substance. Compulsive-like cocaine seeking is usually assessed following an extended free access to the drug (several hours per day), promoting escalation of cocaine intake (*16*), by pairing seeking responses with an aversive footshock (*5*), which identifies 20-30% of individuals as compulsive-like seekers. This behavioral trait is not observed in control animals, which had no extended drug access (*3, 5, 17*), suggesting its necessary contribution to the emergence of compulsive-like seeking. However, such hypothesis cannot be directly tested with the classic procedure, which independently assesses drug intake and compulsive-like seeking, several days apart. Also, the development of compulsion, and its associated mechanisms, cannot be explored since vulnerable individuals display compulsive-like seeking upon their first exposure to the punishment contingency. Thus, when and how does compulsive-like cocaine seeking emerge remains poorly documented (*18*).

### A new behavioral task to concomitantly assess both cocaine intake and seeking

To assess the influence of extended cocaine on the emergence of compulsive-like seeking, we developed a “Punished-Seeking during Extended access” task (PSE), by inserting punished seeking trials within cocaine extended access sessions (**Fig. 1A**). Animals were first trained on a seeking/taking chained schedule of reinforcement (*3*), in which animals have to press a seeking lever under a random interval (1 to 240s). This lever press was never reinforced, but gave access to a second, taking lever, pressing on which triggered a single cocaine infusion (250 μg/90 μL over 5 sec). Following training, animals were then subjected to 15 PSE sessions. Each session started with an initial phase of a 3-hour period of free self-administration of cocaine, under FR1 schedule of reinforcement, followed by a 2-hour seeking phase. Here, half of the completed seeking cycles ended with a mild unpredictable and unescapable electric foot shock (0.5 mA, 0.5 sec), and access to the cocaine lever, to mimic the adverse consequences of the seeking behavior. The daily session ended with another 3-hour period of free access to cocaine.

**Fig. 1.**
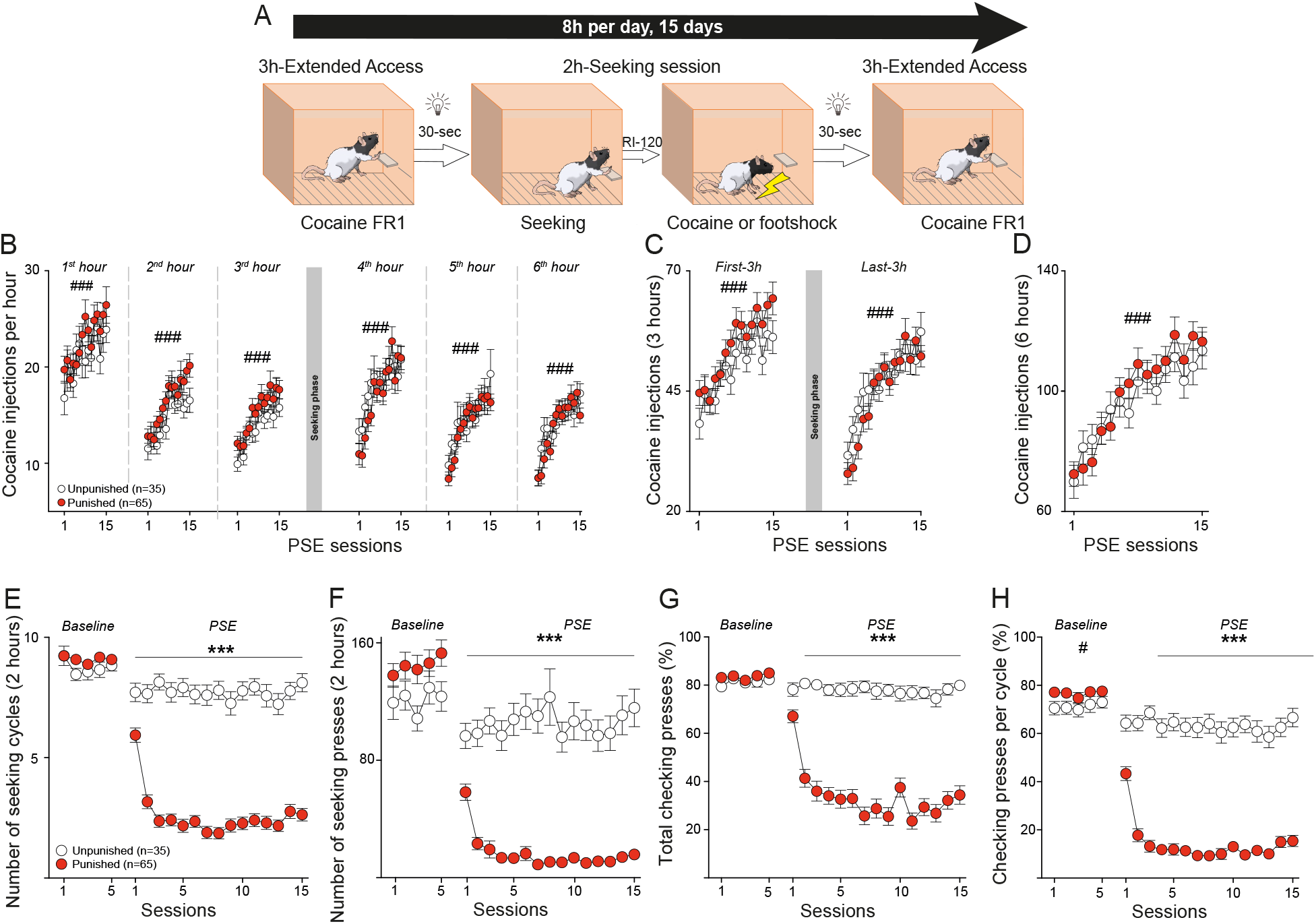
PSE-induced escalation of cocaine and reduction of seeking. (**A**) Schematic of the PSE task: 8h-daily session was run for 15 days (**B**) Hourly cocaine intake PSE-Punished and Unpunished rats (mixed-effects model: *1*^*st*^ *hour*: session effect, *F*_8.848,865.2_ = 5.495, *p* < 0.001; group effect, *F*_1,98_ = 1.421, *n.s*.; *2*^*nd*^ *hour*: session effect, *F*_8.748,855.4_ = 12.57, *p* < 0.001; group effect, *F*_1,98_ = 2.485, *n.s*.; *3*^*rd*^ *hour*: session effect, *F*_7.884,771_ = 12.21, *p* < 0.001; group effect, *F*_1,98_ = 2.003, *n.s*.; *4*^*th*^ *hour*: session effect, *F*_9.803,958.6_ = 11.13, *p* < 0.001; group effect, *F*_1,98_ = 0.5875, *n.s*.; *5*^*th*^ *hour*: session effect, *F*_7.830,765.6_ = 11.02, *p* < 0.001; group effect, *F*_1,98_ = 0.3958, *n.s*.; *6*^*th*^ *hour*: session effect, *F*_8.837,864.1_ = 14.64, *p* < 0.001; group effect, *F*_1,98_ = 0.2184, *n.s*.). (**C**) Cocaine intake per 3-hour blocks (mixed-effects model: *First-3h*: session effect, *F*_7.394,723.1_ = 14,39, *p* < 0.001; group effect, *F*_1,98_ = 2.085, *n.s*.; *Last-3h*: session effect, *F*_8.079,790_ = 17.7, *p* < 0.001; group effect, *F*_1,98_ = 0.4591, *n.s*.). (**D**) Total cocaine intake (mixed-effects model: session effect, *F*_6.514,637_ = 22.61, *p* < 0.001; group effect, *F*_1,98_ = 0.2406, *n.s*.). (**E**) Number of seeking cycle completed during unpunished (*Baseline*) and punished (*PSE*) seeking sessions (mixed-effects model: *Baseline*: group effect, *F*_1,98_ = 3.868, *n.s*.; *PSE*: interaction effect, *F*_14,1370_ = 7.513, *p* < 0.001;). (**F**) Number of seeking lever presses (mixed-effects model: *Baseline*: group effect, *F*_1,98_ = 2.157, *n.s*.; *PSE*: interaction effect, *F*_14,1369_ = 3.97, *p* < 0.001). (**G**) Total percentage of checking lever presses (mixed-effects model: *Baseline*: group effect, *F*_1,98_ = 2.384, *n.s*.; *PSE*: interaction effect, *F*_14,1276_ = 4.62, *p* < 0.001). (**H**) Percentage of checking lever presses per cycle completed (mixed-effects model: *Baseline*: group effect, *F*_1,98_ = 4.34, p = 0.0398, interaction effect: *F*_4,392_ = 0.1532, *n.s*.; *PSE*: interaction effect, *F*_14,1275_ = 6.615, *p* < 0.001). Data are mean ± SEM. ^###^*p* < 0.001 general effect; *, *p* < 0.05, ***, *p* < 0.001 *vs*. Punished.

Unpunished control animals increased their intake during each PSE hour (**Fig. 1B**) and both 3h-phases of drug free access throughout the task (**Fig. 1C**), indicating that adding a seeking phase during extended cocaine access did not occlude the development of escalation of intake (**Fig. 1D**). Likewise, they maintained high seeking performance during the 15 PSE sessions (**Fig. 1E**), evidencing that loss of control over cocaine intake does not impact their willingness to seek a small amount of cocaine (maximum of 10 infusions per 2h-seeking session) despite the subsequent phase of drug free access. In contrast, punishment contingency resulted in a marked reduction of seeking performance, in terms of number of cycles completed and seeking lever presses (**Fig. 1F**). It further decreased checking presses (**Fig. 1, G and H**), those unnecessary to complete a seeking cycle, which assess the animals’ persistent motivation to assess drug availability. Punishment-induced seeking suppression did not however impact cocaine intake, since animals progressively increased their consumption during both phases of extended access, similarly to unpunished individuals.

### PSE induces escalation of compulsive-like cocaine seeking

We next investigated individual’s seeking performances under punishment contingency to identify compulsive rats. Punishment delivery schedule was pseudorandom, so that the difference between both seeking outcomes (punishment or access to cocaine lever) could not exceed two (*3*). As such, animals completing three seeking cycles within a single session experienced both outcomes, highlighting their willingness to seek drug despite the punishment risk. We computed an individual “Compulsivity Score”, which averages the number of seeking cycle completed during the last five sessions of the PSE task (**Fig. 2A**). Animals were then classified as “Compulsive” (Compulsivity Score ≥ 3) or “Non-Compulsive” (**Fig. 2B**), yielding a bimodal distribution of individuals, composed of a normal, and lognormal, distribution for the non-compulsive and compulsive groups, respectively (**Fig. 2C**), as previously reported with the classical sequential model (*2, 3, 5, 19*). Compulsive animals also performed more seeking and checking lever presses during the last five PSE sessions (**Fig. 2, D, E and F**).

**Fig. 2.**
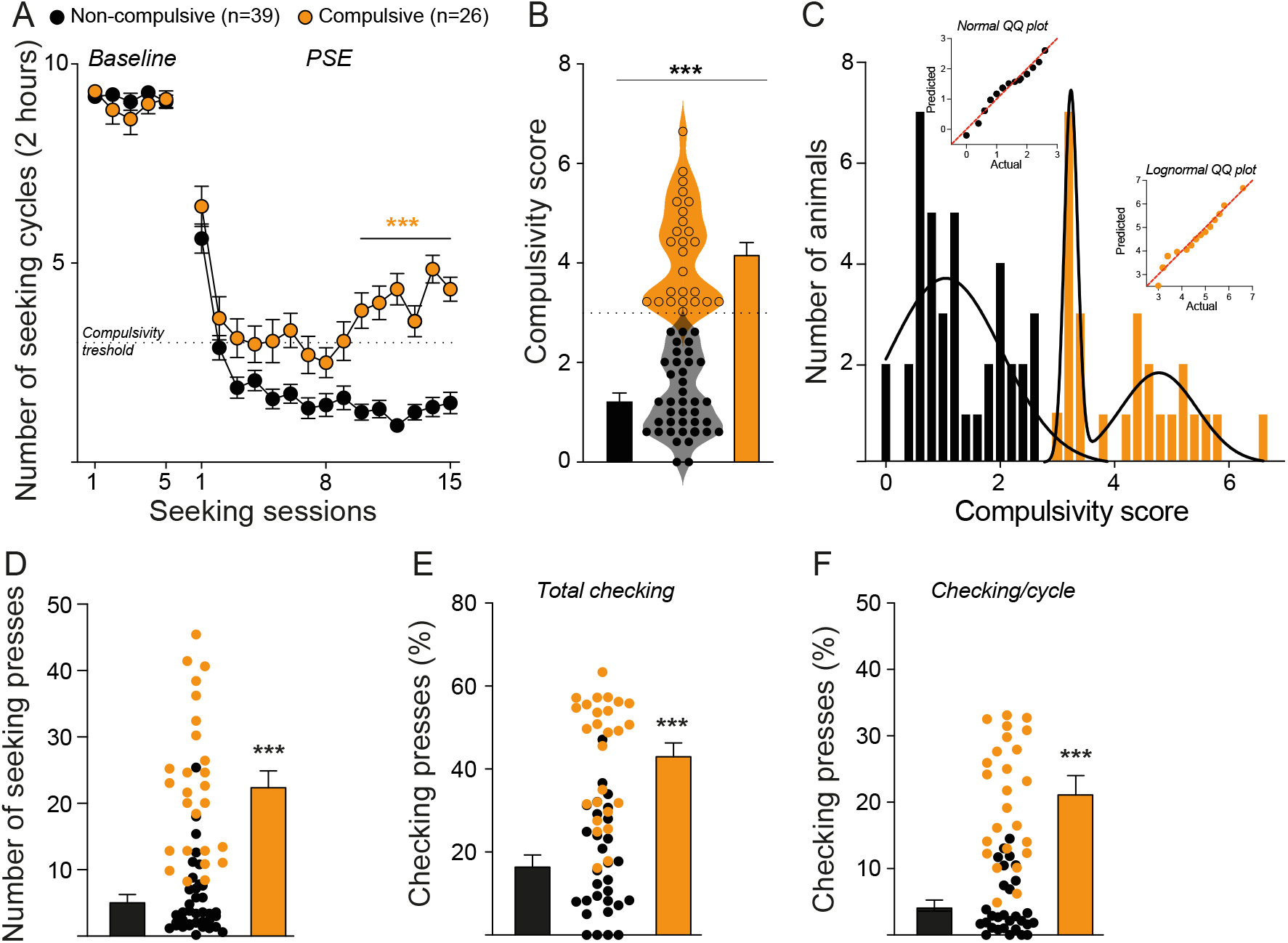
PSE uncovers development of compulsive-like cocaine in a subset of rats. (**A**) Number of seeking cycles completed in compulsive and non-compulsive rats during unpunished (baseline) and punished (PSE) seeking sessions (mixed-effects model: *Baseline*: group effect, *F*_1,63_ = 0.6109, *n.s*.; *PSE*: interaction effect, *F*_14,880_ = 5.027, *p* < 0.001; Šídák post hoc test \*\*\**p* < 0.001 *vs*. Non-compulsive). (**B**) Compulsivity score (Unpaired *t* test: *t*_63_ = 13.43, \*\*\**p* < 0.001). (**C**) Normal and lognormal distribution of and rats, respectively (D’agostino & Pearson test: Non-compulsive: *K2* = 5.187; Compulsive: *K2* = 5.17). (**D**) Number of seeking lever presses during the last five PSE sessions (Unpaired *t* test: *t*_63_ = 8.243, \*\*\**p* < 0.001). (**E**) Percentage of total checking lever presses during the last five PSE sessions (Unpaired *t* test: *t*_54_ = 7.447, \*\*\**p* < 0.001). (**F**) Percentage of checking lever presses per seeking cycle completed during the last five PSE sessions (Unpaired *t* test: *t*_53_ = 8.406, \*\*\**p* < 0.001). Data are mean ± SEM. ***, *p* < 0.001 *vs*. Punished. Dashed lines indicate the compulsivity threshold above which animals are classified as Compulsive.

Both populations displayed similar patterns of cocaine seeking during baseline training and the initial PSE session (**Fig. 3**). Repeated punishment exposure initially reduced seeking performances in both groups, indicating that individuals had learnt the negative consequences of the behavior. Compulsive-like cocaine seeking progressively emerged in some animals (**Fig. 2A, 3A, B and C**), concomitant to a gradual diminution in the latency to initiate seeking during the middle (6-10) and late (11-15) PES sessions (**Fig. 3D**), with no changes in their latency to press the cocaine lever upon availability (**Fig. 3E**). Compulsive animals were also more prone to reinstate cocaine seeking following the first seeking outcome (**Fig. 3F**), regardless of its rewarding nature (cocaine, **Fig. 3G, and H**; shock **Fig. 3I, and J**), evidencing their willingness to reengage in a risky seeking cycle. In contrast, behavioral dynamics of non-compulsive individuals showed a persistent reduction in cocaine seeking behaviors throughout the task.

**Fig. 3.**
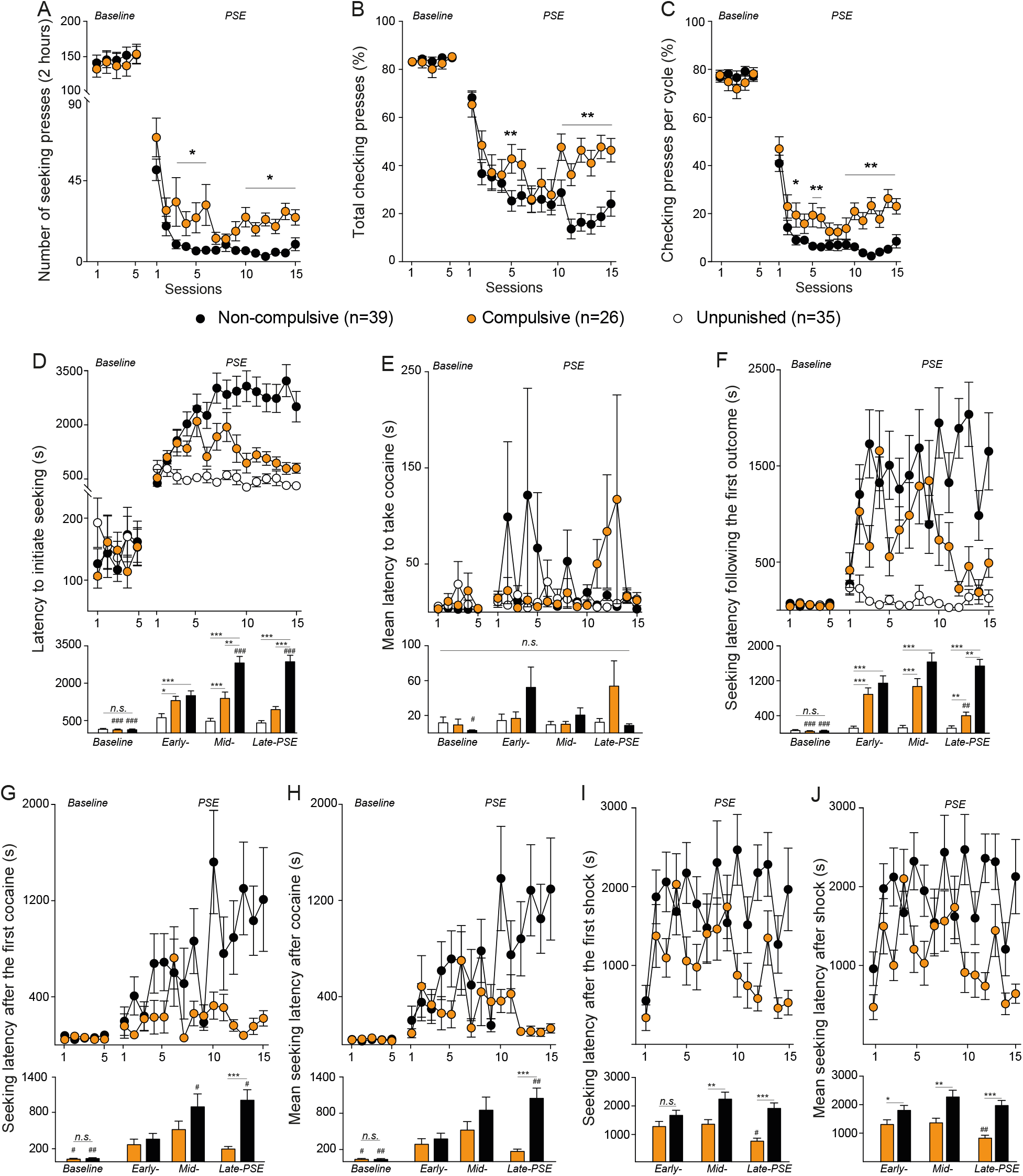
Behavioral features of PSE-induced compulsive-like cocaine seeking. (**A**) Number of seeking lever presses in Non-compulsive and Compulsive rats during unpunished (*Baseline*) and punished (*PSE*) seeking sessions (mixed-effects model: *Baseline*: group effect, *F*_1,63_ = 0.1823, *n.s*.; *PSE*: session effect, *F*_14,879_ = 15.27, *p* < 0.001, group effect, *F*_1,63_ = 29.91, *p* < 0.001). (**B**) Total percentage of checking lever presses (mixed-effects model: *Baseline*: group effect, *F*_1,63_ = 0.5465, *n.s*.; *PSE*: session effect, *F*_14,791_ = 2.57, *p* < 0.01). (**C**) Percentage of checking lever presses per cycle completed (mixed-effects model: *Baseline*: group effect, *F*_1,63_ = 0.5377, *n.s*.; *PSE*: session effect, *F*_14,790_ = 1.78, *p* < 0.05). (**D**) Latency to initiate cocaine seeking in Non-compulsive, Compulsive, and Unpunished rats. Top: time course of responses (mixed-effects model: *Baseline*: group effect, *F*_2,97_ = 0.6805, *n.s*.; *PSE*: interaction effect, *F*_6,291_ = 21.47, *p* < 0.001). Bottom: 5-session blocks analysis (mixed-effects model: interaction effect, *F*_2,97_, *p* < 0.001). (**E**) Mean latency to take cocaine after cycle completion. Top: time course of responses (mixed-effects model: *Baseline*: group effect, *F*_2,97_ = 1.026, *n.s*.; *PSE*: session effect *F*_4.411,303.1_ = 0.6049, *n.s*., group effect, *F*_2,97_ = 1.387, *n.s*.). Bottom: 5-session blocks analysis (mixed-effects model: interaction effect, *F*_6,278_ = 2.642, *p* < 0.05). (**F**) Latency to reinitiate seeking after the first outcome. Top: time course of responses (mixed-effects model: *Baseline*: group effect, *F*_2,97_ = 0.313, *n.s*.; *PSE*: interaction effect *F*_28,1147_ = 2.943, *p* < 0.001). Bottom: 5-session blocks analysis (mixed-effects model: interaction effect, *F*_6,288_ = 13.67, *p* < 0.001). (**G**) Latency to reinitiate seeking after the first cocaine. Top: time course of responses (mixed-effects model: *Baseline*: group effect, *F*_1,63_ = 0.572, *n.s*.; *PSE*: interaction effect *F*_14,465_= 2.37, *p* < 0.01). Bottom: 5-session blocks analysis (mixed-effects model: interaction effect, *F*_2,173_ = 4.616, *p* < 0.01). (**H**) Mean latency to reinitiate seeking after cocaine. Top: time course of responses (mixed-effects model: *Baseline*: group effect, *F*_1,63_ = 0.01463, *n.s*.; *PSE*: interaction effect *F*_14,478_ = 2.705, *p* < 0.001). Bottom: 5-session blocks analysis (mixed-effects model: interaction effect, *F*_3,173_ = 5.591, *p* < 0.01). (**I**) Mean latency to reinitiate seeking after the first shock. Top: time course of responses (mixed-effects model: *PSE*: session effect *F*_14,595_ = 3.323, *p* < 0.001, group effect, *F*_1,63_ = 20.71, *p* < 0.001). Bottom: 5-session blocks analysis (mixed-effects model: session effect, *F*_1.883,113.9_ = 4.182, *p* < 0.05, group effect, *F*_1,63_ = 19.85, *p* < 0.001). (**J**) Mean latency to reinitiate seeking after shocks. Top: time course of responses (mixed-effects model: *PSE*: interaction effect, *F*_14,598_ = 1.904, *p* < 0.05). Bottom: 5-session blocks analysis (mixed-effects model: session effect, *F*_1.880,113.8_ = 3.931, *p* < 0.05, group effect, *F*_1,63_ = 23.93, *p* < 0.001). Šídák post hoc test ^#^p < 0.05, _##_p < 0.01, ^###^p < 0.001 *vs. Early*; \**p* < 0.05, \*\**p* < 0.01, \*\*\**p* < 0.001 *vs*. Non-compulsive or Unpunished. Data are mean ± SEM.

Compulsive-like seeking behaviors is usually assessed with a sequential approach, in which individuals’ resistance to punishment is tested after, i.e. independently of, an extended period of cocaine free access (*2, 3, 5, 6, 14, 17*). When subjected to post-PSE punishment seeking sessions, with no extended access to cocaine (**Fig 4A**), compulsive animals still displayed pathological patterns of drug seeking **(Fig. 4B**), highlighting the persistence of PSE-induced compulsive cocaine seeking. Additionally, when exposed to punishment contingency after the PSE task, a subset of control rats that did not experience punishment during PSE, immediately expressed compulsive-like cocaine seeking. This suggests that compulsive seeking can also be developed in the absence of punishment, which however did not occlude its subsequent expression when facing negative consequences of the behavior.

**Fig. 4.**
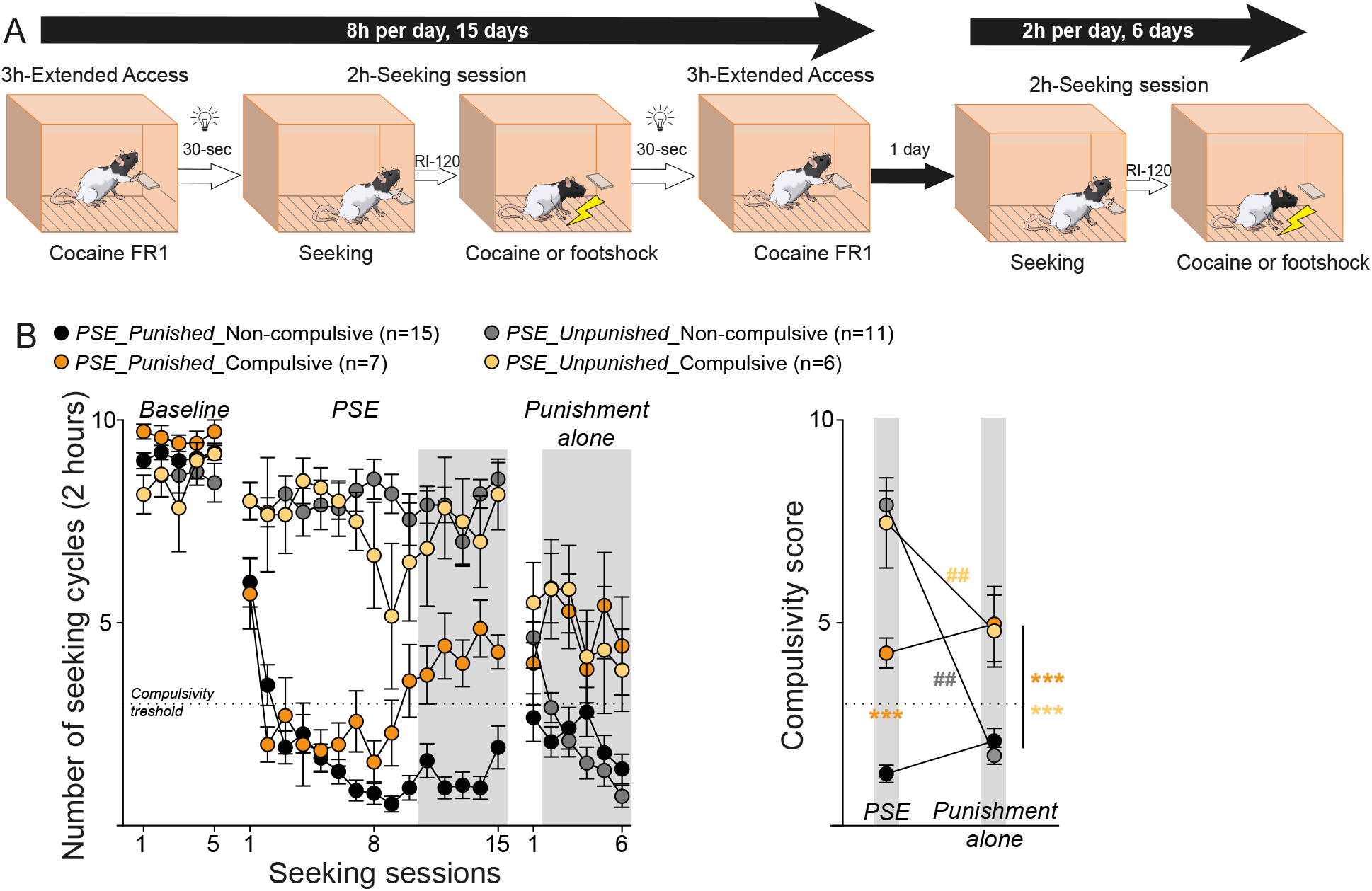
PSE-induced expression of compulsive-like seeking is independent of cocaine extended access. (**A**) Schematic of the procedure: following PSE task (8 hours per day for 15 days), punished seeking (no extended access to cocaine) was assessed for 6 days (2 hours per day). (**B**) *Left*: Number of seeking cycle completed in Compulsive and Non-compulsive PSE_Punished, and Compulsive and Non-compulsive PSE_Unpunished rats during *Baseline* (no punishment), *PSE* (± punishment), and *Punishment alone* (no extended access) sessions (mixed-effects model: *PSE_Unpunished*_*Baseline*: group effect, *F*_1,15_ = 0.322, *n.s*.; *PSE*: interaction effect *F*_1,15_ = 0.5203, *n.s*.; *Punishment alone*: session effect: *F*_5,75_ = 5.627, *p* < 0.001; group effect: *F*_1,15_ = 16.97, *p* < 0.001; *PSE_Punished*_*Baseline*: group effect, *F*_1,20_ = 4.932, *p* < 0.05; session effect *F*_4,80_ = 0.3684, *n.s*.; interaction effect *F*_4,80_ = 0.1937, *n.s*.; *PSE*: interaction effect *F*_14,280_ = 5.025, p < 0.001; *Punishment alone*: interaction effect: *F*_5,100_ = 2.373, *p* < 0.05). Grey rectangles indicate the sessions used to establish the compulsivity score in *Right*: (sessions effect, *F*_1,20_ = 4.533, *p* = 0.05, group effect: *F*_1,20_ = 34.92, *p* < 0.001; Šídák post hoc test ^##^*p* < 0.01 *vs*. PSE; \*\*\**p* < 0.001 *vs*. Non-compulsive). Data are mean ± SEM. Dashed lines indicate the compulsivity threshold above which animals are classified as Compulsive.

### Compulsive-like cocaine seeking emerges following escalation of intake

Like unpunished animals, PSE-punished rats increased their hourly cocaine consumption during both 3-hour phases of extended access (**Fig. 5A, and B**), resulting in a similar escalation of cocaine intake (**Fig. 5C**) in both Compulsive and Non-compulsive rats, which did not impact their promptness to engage cocaine consumption (**Fig. 5D**). Thorough examination of the seeking time course behavioral dynamics revealed that compulsion started to emerge after eight PSE sessions, a behavioral time point at which animals have already escalated their intake. This suggests that prolonged extended access to cocaine may gate the emergence of compulsive seeking in vulnerable individuals. We thus split the PSE task to investigate intake dynamics during the early (sessions 1 to 8) and the late (sessions 9 to 15) phases of the task. Linear regression of cocaine intake data revealed different dynamics of consumption between the beginning and the end of the extended access periods. While both compulsive and non-compulsive rats displayed similar intake dynamics throughout the first 3h-period (**Fig. 5E**), only non-compulsive rats kept increasing their intake until the end of the protocol. In contrast, intake dynamics of compulsive rats revealed that they ceased to increase their intake after 8 PSE sessions, reaching an ‘escalated intake’ *plateau*. Comparison of global taking and seeking dynamics during PSE showed that stabilization of escalated cocaine intake during the late extended access periods parallels the emergence of pathological cocaine seeking in compulsive rats (**Fig. 1F**). This highlights the necessary role of extended and prolonged cocaine intake for the subsequent development of compulsive cocaine seeking (*3, 5, 17*). To causally confirm this gating role, the length of the extended access periods was reduced from 3 to 1 hour, to limit cocaine availability and consumption. Although punishment exposure induced a rapid decrease in cocaine intake, no escalation of cocaine intake was observed in punished and unpunished rats, which exhibited stable consumption throughout the short PSE task (**Fig. 5G, and H**). In turn, none of the PSE-punished animals developed compulsive-like cocaine seeking (**Fig. 5I, and J**), confirming that pathological compulsive seeking can only emerge in a pre-existing cocaine escalated state. Thus, emergence of compulsive seeking requires an extended access to cocaine over a prolonged period, but once developed, its expression is independent of drug availability.

**Fig. 5.**
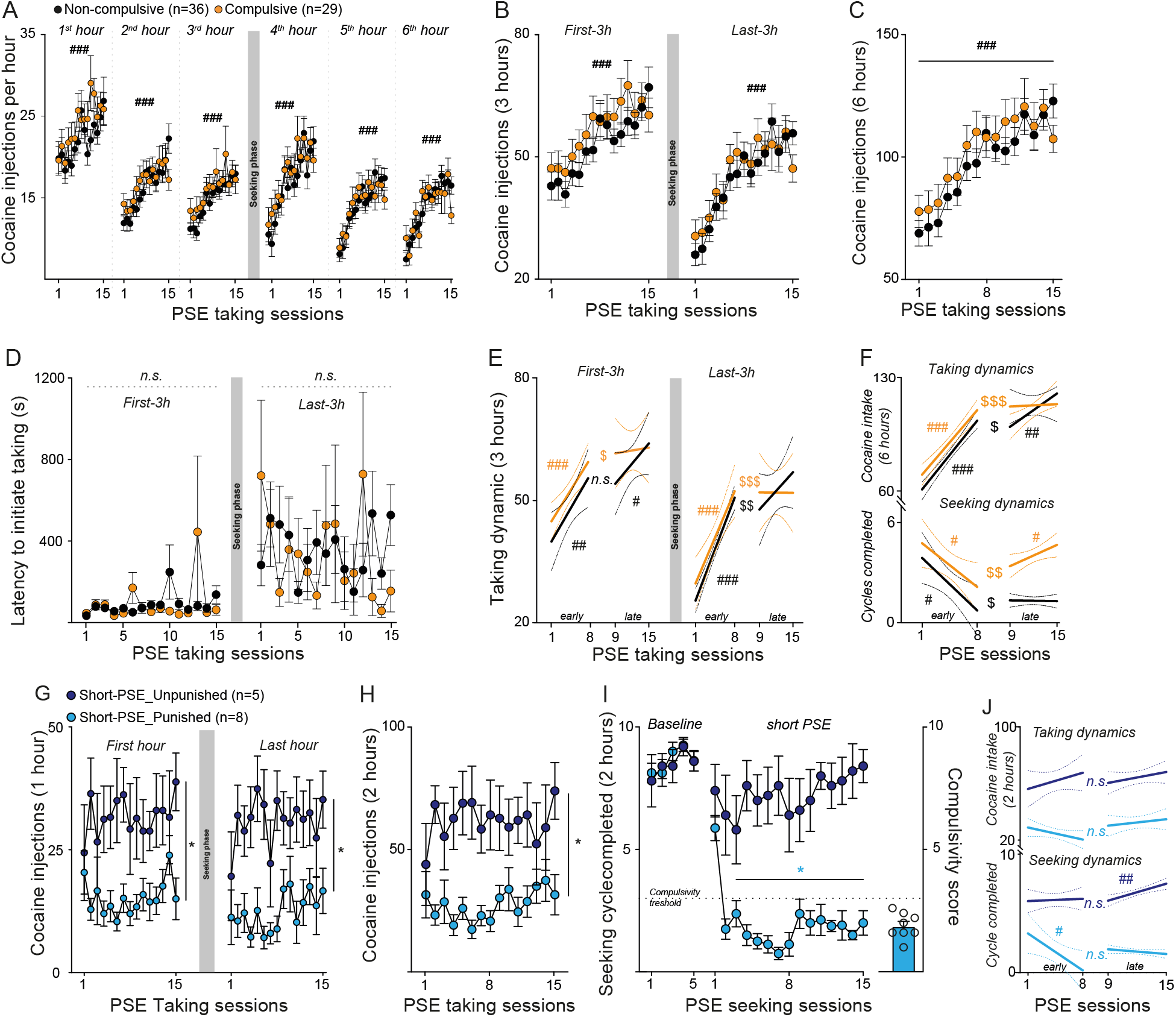
Extended access to cocaine gates it compulsive-like seeking. (**A**) Hourly cocaine intake of compulsive and non-compulsive rats (mixed-effects model: *1*^*st*^ *hour*: session effect, *F*_7.784,488.7_ = 4.462, *p* < 0.001; group effect, *F*_1,63_ = 1.553, *n.s*.; *2*^*nd*^ *hour*: session effect, *F*_7.623,478.6_= 9.132, *p* < 0.001; group effect, *F*_1,63_ = 0.2623, *n.s*.; *3*^*rd*^ *hour*: session effect, *F*_7.731,485.4_ = 8.064, *p* < 0.001; group effect, *F*_1,63_ = 1.162, *n.s*.; *4*^*th*^ *hour*: session effect, *F*_9.646,605.7_ = 10.46, *p* < 0.001; group effect, *F*_1,63_ = 0.9032, *n.s*.; *5*^*th*^ *hour*: session effect, *F*_9.106,571.1_ = 9.874, *p* < 0.001; group effect, *F*_1,63_ = 0.05764, *n.s*.; *6*^*th*^ *hour*: session effect, *F*_8.868,556.8_ = 11.34, *p* < 0.001; group effect, *F*_1,63_ = 0.01753, *n.s*.). (**B**) Cocaine intake per 3-hour blocks (mixed-effects model: *First-3h*: session effect, *F*_6.947,436.2_ = 10.07, *p* < 0.001; group effect, *F*_1,63_ = 1.004, *n.s*.; *Last-3h*: session effect, *F*_8.517,534.7_ = 14.24, *p* < 0.001; group effect, *F*_1,63_ = 0.2004, *n.s*.). (**C**) Total cocaine intake (mixed-effects model: session effect, *F*_6.936,436_ = 16.77, *p* < 0.001; group effect, *F*_1,63_ = 0.5976, *n.s*.). (**D**) Latency to initiate cocaine taking of Non-compulsive and Compulsive rats (mixed-effects model: *First-3h*: group effect, *F*_1,63_= 0.02197, *n.s*.; *Last-3h*: group effect, *F*_1,63_= 0.05724). (**E**) Cocaine taking behavioral dynamics between the *early* (sessions 1 to 8) and *late* (sessions 9 to 15) PSE sessions (linear regression: first-3h: Non-compulsive *early R*^2^ = 0.7945, *p* = 0.003, *late R*^2^ = 0.6614, *p* = 0.026; Compulsive *early R*^2^ = 0.8904, *p* < 0.001, *late R*^2^ = 0,03014, *n.s*.; last-3h: Non-compulsive *early R*^2^ = 0.9833, *p* < 0.001; *late R*^2^ = 0.5568, *p* = 0.054; Compulsive *early R*^2^ = 0.9099, *p* < 0.001; *late R*^2^ = 0.0005114, *n.s*.; ANCOVA: First-3h *early vs. late*: Non-compulsive: *F*_1,11_ = 0.6212, *n.s*.; Compulsive *F*_1,11_= 8.673, *p* = 0.013; Last-3h *early vs. late*: Non-compulsive: *F*_1,11_ 12.7, *p* = 0.004; Compulsive *F*_1,11_= 18.36, *p* = 0.001). (**F**) Cocaine taking (upper) and seeking (bottom) behavioral dynamics between the *early* (sessions 1 to 8) and *late* (sessions 9 to 15) PSE sessions (linear regression: *early* taking Non-compulsive *R*^2^ = 0.958, *p* < 0.001; Compulsive *R*^2^ = 0.923, *p* < 0.001; *late* taking Non-compulsive *R*^2^ = 0.7686, *p* = 0.01; Compulsive *R*^2^ = 0.008288, *n.s*.; *early* seeking Non-compulsive *R*^2^ = 0.6041, *p* = 0.023; Compulsive *R*^2^ = 0.5401, *p* = 0.038; *late* seeking Non-compulsive *R*^2^ = 0.02649, *n.s*.; Compulsive *R*^2^ = 0.6045, *p* = 0.04; ANCOVA: *early vs. late* taking: Non-compulsive: *F*_1,11_ = 8.586, *p* = 0.014; Compulsive: *F*_1,11_ = 19, *p* = 0.001; *early vs. late* seeking: Non-compulsive: *F*_1,11_ = 6.18, *p* = 0.03; Compulsive: *F*_1,11_ = 10.94, *p* = 0.007. (**G**) Cocaine intake during the first and the last hour of the short-PSE task in Punished and Unpunished rats (per 3-hour blocks (mixed-effects model: *First hour*: session effect, *F*_14,153_ = 0.4317, *n.s*.; group effect, *F*_1,11_ = 15.65, *p* < 0.01; *Last hour*: session effect, *F*_14,153_ = 1.116, *n.s*.; group effect, *F*_1,11_ = 16.21, *p* < 0.01). (**H**) Total cocaine intake in short-PES_Unpunished and _Punished rats (mixed-effects model: session effect, *F*_14,153_ = 0.5766, *n.s*.; group effect, *F*_1,11_ = 16.3, *p* < 0.01). (**I**) *left*: Number of seeking cycle completed by short-PES_Unpunished and _Punished rats during *Baseline* (no punishment) and *short-PSE* (± punishment) sessions (mixed-effects model: *Baseline*: group effect, *F*_1,11_ = 0.08633, *n.s*.; *short-PSE*: session effect, *F*_4.865,53.52_ = 0.0916, *n.s*.; group effect, *F*_1,11_ = 108.8 < 0.001). *Right*: Compulsivity score of Short-PSE_Punished rats. (**J**) Cocaine taking (upper) and seeking (bottom) behavioral dynamics between the *early* (sessions 1 to 8) and *late* (sessions 9 to 15) PSE sessions in Short_PSE_Unpunished and _Punished rats (linear regression: *early* taking Unpunished *R*^2^ = 0.219, *n.s*.; Punished *R*^2^ = 0.3958, *n.s*.; *late* taking Unpunished *R*^2^ = 0.3093, *n.s*.; Punished *R*^2^ = 0.1502, *n.s*.; *early* seeking Unpunished *R*^2^ = 0.01326, *n.s*.; Punished *R*^2^ = 0.5558, *p* = 0.034; *late* seeking Unpunished *R*^2^ = 0.8131, *p* < 0.01; Punished *R*^2^ = 0.3686, *n.s*.; ANCOVA: *early vs. late* taking: Unpunished *F*_1,11_ = 0.04226, *n.s*.; Punished: *F*_1,11_ = 3.875, *n.s*.; *early vs. late* seeking: Unpunished: *F*_1,11_ = 3.081, *n.s*.; Punished: *F*_1,11_ = 3.844, *n.s*.). ^#^*p* < 0.05, ^##^*p* < 0.01, ^###^*p* < 0.001; ^$^*p* < 0.05, ^$$^*p* < 0.01, ^$$$^*p* < 0.001 *early vs. late*; \**p* < 0.05 *vs*. Unpunished). Data are mean ± SEM (panels A, B, C, D, G, H, I) or 95% confidence interval (panels E, F, J). The dashed line indicates the compulsivity threshold above which animals are classified as Compulsive.

## Discussion

Rodent models assessing pathological cocaine seeking after extended period of drug free access hardly recreate the human condition, in which dysregulation of consumption and compulsive pattern of drug seeking and taking likely co-occur. Furthermore, they do not offer the possibility to exactly determine how and when compulsion emerges, as vulnerable individuals promptly exhibit compulsive-like seeking upon their first exposure to punishment (*14, 18*). The PSE task used in the present study circumvents these caveats since pathological seeking is tested within daily periods of free access to the drug, allowing the concomitant assessment of dysregulation of cocaine intake and its compulsive seeking, two important features for addiction diagnosis in human *(1)*. Our results are in line with key features of classical addiction-like models, providing high face validity of the PSE task: (*i*) extended access to cocaine induces a progressive increase in drug consumption, reflecting escalation (i.e. dysregulation) of cocaine intake (*16*); (ii) a subset of animals displays compulsive pattern of cocaine seeking when exposed to punishment (*3, 5*). Importantly, combining punished sessions with extended access periods has no occluding effects on seeking and taking behaviors. Punishment-induced devaluation of the seeking outcome during PSE does not prevent subsequent cocaine taking, since punished and unpunished animals displayed similar levels of cocaine consumption, and escalated their intake during the second 3h-phase of the daily sessions. Similarly, unlimited access to cocaine during the first 3h-phase of free extended access to cocaine did not dampen seeking performances of PSE-unpunished rats.

As expected, punishment contingency induced a general and rapid reduction in animals’ drug seeking behavior. However, while seeking performances of most individuals remained low throughout the PSE task, a subset of animals progressively exhibited pathological seeking pattern, which persisted after PSE, in the absence of extended access to cocaine. Both compulsive and non-compulsive rats displayed similar levels of cocaine intake during extended access phases, with however different dynamics during the last half of PSE. While non-compulsive animals showed a constant increase in intake dynamic throughout the task, compulsive rats showed an abrupt shift in their intake toward a persistent steady-state plateau after eight PSE sessions. Strikingly, the stabilization of cocaine *escalated* intake occurred in parallel with the emergence of compulsive-like seeking. This suggests that the development of compulsion requires a certain level of drug intake, obtained during extended access periods. In line with such hypothesis, no compulsive-like seeking was observed in animals with a limited access to cocaine.

The PSE task thus allows the study of the concomitant development of pathological cocaine seeking and taking behaviors, and their relationship. In fact, we observed a sequential escalation-like increase in both drug intake and seeking in compulsive-like animals, with the former slightly preceding the latter, positioning dysregulation of cocaine intake as a permissive step for the subsequent development of compulsive seeking in vulnerable individuals. This task thus offers a unique opportunity to investigate mechanisms sustaining the emergence of compulsion, which remain largely unknown. Spreading its use should improve our understanding of brain processes at stack, eventually leading to new therapeutic strategies to reduce or prevent the development the drug seeking phase to avoid extended access to the drug. Ahead of this, our results should advocate for the necessity to develop prevention policy strategies aiming at limiting extended and prolonged access to a substance to prevent the emergence of subsequent pathological pattern of drug seeking and use.

## Acknowledgments

We thank the National Institute on Drug Abuse Drug Supply Program (NDSP) for generously providing cocaine. We thank David Meunier for elaboration of the python pipeline for extraction of behavioral data, and the institute’s platforms involved (PAG, NIT, S-prim, UAR 2018). No editing services or AI-assisted technologies were used in preparation of the manuscript.

## Funding

Agence National de la Recherche JCJC_ANR-21-CE16-0002-01 (MD)

NARSAD Young Investigator Grant from the Brain & Behavioral Research

Foundation_300015 (MD)

Fondation NRJ_2019 (CB)

## Author contributions

Conceptualization: MTP, MD

Methodology: MTP, CB, JMG, MD

Investigation: MTP, JLdP, LV, MW, MD

Visualization: MTP, MD

Data curation: MTP, JLdP, JMG, MD

Funding acquisition: CB, MD

Project administration: MD

Supervision: MD

Writing – original draft: MTP, MD

Writing – review & editing: MTP, JLdP, LV, MW, CB, JMG, MD

## Data and materials availability

All data are available in the main text or the supplementary materials.

## Materials and Methods

### Animals

Adult Lister Hooded males (∼4-8week-old, Charles River, N = 123) were paired housed, in Plexiglas cages and maintained on an inverted 12h light/dark cycle (light onset at 7.30 p.m.) with food and water available ad libitum, in a temperature- and humidity-controlled environment. Experimental manipulations were carried out in the animals’ dark cycle between 8:00 a.m. and 6:00 p.m. All handling and experimental procedures complied with the recommendations for animal experiments issued by the European Commission directives 219/1990, 220/1990, and 2010/63, and approved by Ethic Committee APAFIS# #37070-202204201748941 v3/ #43676-2023060614092748 v4).

### Surgery

Rats were implanted with a chronic indwelling intravenous catheter. Rats were anesthetized with ketamine (Imalgen, Merial, 100 mg/kg, s.c.) and medetominine (Domitor, Janssen, 30 mg/kg, s.c.) following a preventive long-acting antibiotic treatment (amoxicillin, Duphamox LA, Pfizer, 100 mg/kg, s.c.). The homemade silicone catheter (0.012-inch inside diameter, 0.025-inch outside diameter) was inserted and secured into the right jugular vein. The other extremity of the catheter was placed subcutaneously in the mid-scapular region and connected to a guide cannula secured with dental cement. After surgery, rats were awakened with an injection of atipamezol (Antisedan, Janssen, 0.15 mg/kg i.m.) and allowed to recover for at least 5 days with ad libitum access to food and water. The catheters were daily flushed during the recovery period, and just before and after each self-administration session with a saline solution containing heparin (Sanofi, 3 g/l) and gentamicine (Pangram® 4%, Virbac) to maintain their patency and to prevent infection. Catheters were also regularly tested with propofol (Propovet, Abbott, 10 mg/ml) to confirm their patency.

### Behavioral apparatus

Behavioral experiments were performed during animals’ dark phase, and took place in standard rat operant chambers (MedAssociates), located in sound-attenuating cubicles, equipped with a house light, two retractable levers set 7 cm above the metallic grid floor through which an electric foot shock could be delivered via a generator (MedAssociates). A cue light was positioned 8 cm above each lever. For each rat, one lever was randomly paired with cocaine infusion (taking-lever) while the other one was designated as the seeking-lever. For intravenous drug administration, the stainless-steel guide cannula of the catheter was connected through steel-protected tygon tubing to a swivel (Plastics One) and then an infusion pump (MedAssociates). Data were acquired on a PC running MED-PC IV-V (MedAssociates). Sessions lasted for 2h, 4h or 8h (see below for detailed procedures).

### ‘Punished Seeking during Extended access’ task

After recovery from surgery, rats began cocaine self-administration training using the seeking-taking chained schedule. Self-administration training was divided into three distinct phases: acquisition of the taking response, training on the seeking-taking chained, PSE.

#### Acquisition of the taking response

Each trial started with the illumination of the house light and the insertion of the taking-lever. One press on the lever, under a fixed ratio schedule (FR1), resulted in the delivery of a single infusion of cocaine (250 μg/90 μL over 5 seconds, (National Institute on Drug Abuse Drug Supply Program). Cocaine infusions were paired with illumination of the cue light (5 seconds) above the taking lever, retraction of the taking-lever, and extinction of the house light for 5 seconds. Following a 20 second-time out-period (TO), another trial was initiated with the insertion of the taking-lever. Training of the taking response continued, typically for four to eight sessions, until animals reached a stable level of cocaine.

#### Training on the seeking-taking chain schedule

Each cycle started with the illumination of the house light, insertion of the seeking-lever. A single press on the seeking-lever initiated a random interval (RI < schedule of 2 seconds (0.1 < RI < 4seconds). Seeking-lever presses within the RI (unnecessary seeking lever presses) were recorded as checking lever presses (see below). The first seeking-lever press following the end of the RI, ending the seeking-taking cycle, triggered its retraction, and the insertion of the taking-lever. Press on that lever delivered the cocaine administration, paired with the illumination of the associated cue-light, followed by a 20-second TO, during which both levers were retracted.

Following TO, seeking lever was re-inserted to begin the next seek-take cycle. With acquisition of the seeking-taking responses, RI and TO were progressively increase to reach 20, 60, 120 seconds, and 1, 5, 10 minutes, respectively. At the end of training under the seeking-taking schedule, animals were allowed to complete up to 10 seeking cycles during each 2h-session of the RI120-TO10 schedule.

#### PSE

Following training (ending with at least five RI120-10TO baseline sessions), all animals were subjected to the PSE task. Each daily sessions started with the illumination of the house light and the insertion of the taking-lever. Each press on this lever triggered cocaine delivery and the illumination of the associated cue-light, followed by a 20s-TO. During the first phase of PSE (Extended access 1), animals can freely and unlimitedly self-administer cocaine for 3 hours (1 hour for the short version of the task). The end of this was signaled by the retraction of the seeking lever, and extinction of the house light for 15 seconds.

Following a 15-second period of house light flashing, the beginning of the second PSE phase (Seeking session) is initiated with the illumination of the house light and the insertion of the seeking lever only. Here, animals can seek cocaine under a RI120-TO10 schedule for 2 hours.

Half of the seeking cycle completed resulted in mild foot shock delivery (0.5 mA, 0.5 s) to the animals’ paws, with no access to the taking lever. Punishment was administered following the first press on the seeking lever after the RI120 schedule has elapsed. The other half of the RI120 completed triggered the insertion of the taking lever, which press initiated cocaine delivery and illumination of the associated cue-light. Seeking cycle’s outcomes (foot-shock or insertion of the cocaine taking lever) were delivered in a pseudorandomized manner, so that difference between both outcomes cannot exceed 2. As such, if the two first seeking cycles lead to two consecutive foot shocks, the third cycle will automatically give access to the cocaine-taking lever. Thus, depending on their performances on the punished seeking-taking task, animals can complete up to 10 cycles, leading to a maximum of five foot-shocks and five cocaine infusions. The end of the seeking session was signaled by retraction of the seeking lever, if inserted, and the extinction of the house light for 15 seconds.

Following a 15-second period of house light flashing, the third phase of PSE (Extended access 2) was initiated. As in Extended access 1 period, animals can freely and unlimitedly self-administer cocaine for 1 or 3 hours.

### Behavioral classification

As previously described (*3*), compulsivity score was computed from the number of seeking cycle completed. Here, we averaged the number of completed cycles (regardless of their outcomes) during the last five sessions of PSE. Animals with a compulsivity score ≥ 3 (i.e., more than 30% of the 10-daily seeking cycles during the last five sessions were completed) were classified as ‘compulsive’. Consequently, animals performing more than 3 cycles per session were necessarily exposed to both outcomes. To further appreciate the compulsivity of the animals, we also computed the total percentage of checking presses (those within the RI, with no behavioral consequences), and the percentage of checking responses per cycle completed, as follow: 100* ((checking lever presses/total seeking lever presses) * (cycles completed/total cycles)).

### Statistical analyses

Data are expressed as mean ± SEM, or 95% confidence interval for regression analyzes, with the exact sample size indicated for each group in the figures. Statiostics are indicated in figure legends. Data were extracted using a custom-made python pipeline, and analyzed with two-tailed *t* test, mixed two-ANOVA, mixed-effect models (REML, in case of absence of data, *e.g*. no seeking presses during seeking sessions), regression, and ANCOVA, followed by a Šídák post hoc test when appropriate, using Prism 10 (GraphPad). Only *p*-values ≤ 0.05 were considered significant.

